# A taxonomy based on acoustic features of some Iranian cicadas and calling song description of *Chloropsalta smaragdula* Haupt, 1920 (Hem: Cicadidae) from Isfahan, Iran

**DOI:** 10.1101/2024.01.28.577653

**Authors:** Maedeh Mehdipour, Klaus Riede, Jalal Jalali Sendi, Hossein Zamanian, Akbar Mohammadi-Mobarakeh

## Abstract

This paper compiles parameters of calling songs from 14 cicada species inhabiting Iran. In addition, calling song parameters of *Chloropsalta smaragdula* were analyzed for the first time. A taxonomy based on song parameters was designed, including Iranian cicadas *Tibicen plebejus*, *Cicadatra lorestanica*, *Cicadivetta tibialis*, *Tettigetta golestani*, *Pagiphora annulata*, *Tibicina haematodes*, *Cicada orni, Pagiphora annulata*, *Chloropsalta smaragdula*, *Cicadatra hyalina*, *Psalmocharias querula*, *Cicadatra persica*, *Cicadatra alhageos*, *Cicadatra atra* and *Cicadatra barbodi* out of 44 species reported from Iran. In addition to common acoustic parameters, four new complex spectral features such as variance, kurtosis, spectral centroid and short time energy were used. These additional features were necessary to construct a comprehensive identification key based on acoustic parameters. Our acoustic identification system provides a highly accurate species recognition method, which could be of general relevance in cicada taxonomy.

## Introduction

In this study, published calling songs of 15 out of 44 species of cicadas from Iran were re-described using spectral and temporal paramaters, together with four complex features of calling songs: variance, kurtosis, spectral centroid and short time energy. In addition, recordings of the calling song of *Chloropsalta smaragdula* (Subfamily Cicadinae: Tribe Gaeanini) were analyzed for the first time.

Cicadas produce complex acoustic signals which are species-specific but differ according to biological context and behavior (e.g. Claridge, 1985; Boulard, 1995; Quartau, 1995; Simões et al., 2000). Males produce a “calling song” to attract females at long range. These constitute the first step in pair formation. In many species singing males switch from calling song to “courtship signal” as soon as the female joins the male before mating (Lei et al. 1994; Chou et al. 1997; Cooley and Marshall 2001). Calling and courtship songs are important for mate recognition and necessary for a successful reproduction (Claridge 1985; Marshall and Cooley 2000; Sueur and Aubin 2003; Tishechkin 2003). “Rivalry signals” are produced in cicada male–male interactions at close quarters to maintain distance among competing males (Popov 1975; Cocroft and Pogue 1996; Tishechkin 2003). In some species males produce “distress signals” in dangerous situations, e.g. when stressed by predators (Tishechkin 2003). Females are silent, but some species generate short wing clicks as an acceptance signal (Cooley and Marshall 2001).

For each species songs can be characterized by temporal patterns and spectral features. Boulard (2004) used these acoustic features to create “Cards for Identification by Acoustics”, and introduced the terms sonotypes and etho-sonotypes to differentiate between sibling or cryptic species. (e.g. Dugdale and Fleming, 1978; Clardige 1985; Tishechkin 2003; Gogala and Trilar 2004; Quartau and Simões, 2006; Puissant and Sueur, 2010; Marshall et al. 2011). Therefore, song parameters can be used for species differentiation and description, showing remarkable differences even among closely related species (e.g. Alexander and Moore 1962; Villet 1988, 1989; Boulard 1995; Marshall and Cooley 2000; Sanborn and Phillips 2001; Ewart 2005)

Since they are important for reproductive isolation, songs are highly species-specific and can be used to differentiate distinct species and classify their taxonomic status (see Gogala and Trilar (2004) for *Cicadetta montana* sensu lato complex). Ohya (2004) identified 12 species of genus *Tibicen* by 3 features of their calling song: peak frequency, mean frequency and number of pulses per second. Puissant and Sueur (2010) classified 23 species of West European Cicadettini based on acoustic and morphological identification. All these examples demonstrate the importance of bioacoustics for the study of cicada behaviour and taxonomy.

In this work we tried to classify all species of cicada inhabiting Iran by visible spectral and temporal signal features (VSTF). Most VSTF features could be extracted from the literature. For 6 species (*Psalmocharias querula*, *Cicadatra alhageos*, *Cicadatra atra*, *Cicadatra barbodi* and *Cicadatra persica*) VSTF parameter information in papers is insufficient. Therefore, we added mathematical methods based on four complex spectral features (CSF) such as Variance, Kurtosis, Spectral centroid and Short time energy, which allowed reliable classification independent of the signal section. These alternative acoustic identification features demonstrated high accuracy of recognizing species and taxonomy of cicada by acoustic traits.

## Material and Methods

### Acoustical taxonomy

The visible spectral and temporal signal features (VSTF) of calling songs are defined based on the terminology generally used in cicada bioacoustics (e.g. Sueur and Aubin, 2004; Boulard and Eliopolous 2006; Eliopoulos 2006; Pussiant and Sueur 2010). Calling songs are repeated over long periods of time. The repeated element is sometimes called a phrase. Each repeatable phrase usually consists of two subgroups with strong amplitude variations (Fig 1, 2, 3), followed by a silent interphrase gap (Fig 2, 4e). The two subgroups of the phrase are referred to as low amplitude and high amplitude parts (Sueur and Aubin 2004). In the temporal domain, there is big variation of phrase duration, but for most acoustic signals the phrases last for periods of seconds. Each phrase subgroup consists of echemes with a temporal length varying widely between hundredths of seconds down to tenths of milliseconds (Fig 2, 3, 4, 5, 6, 7). Each echeme consists of syllables in the millisecond range. Syllables consist of pure pulses, representing the basic element of sound or vibration production, of the order of tenths of milliseconds (Fig 4e, d, f) (Eliopoulos, 2006). In the frequency domain, the dominant frequency is the frequency of the maximum amplitude on the spectrogram (Fig 8,9) (Boulard and Eliopolous 2006; Pussiant and Sueur 2010).

**Figure 1:**
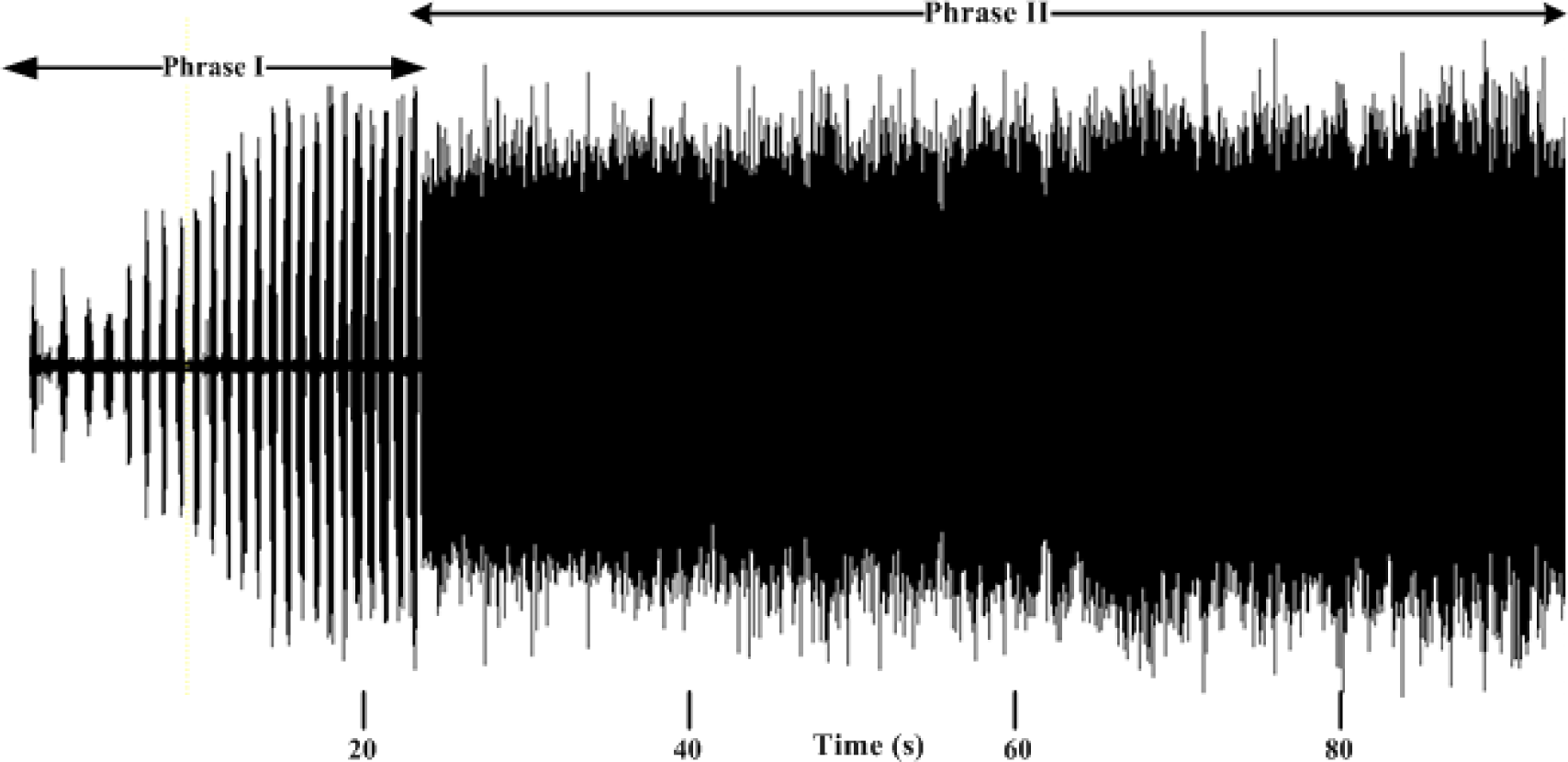
an example of two type of phrase; calling song of *Chloropsalta smaragdula*

**Figure 2:**
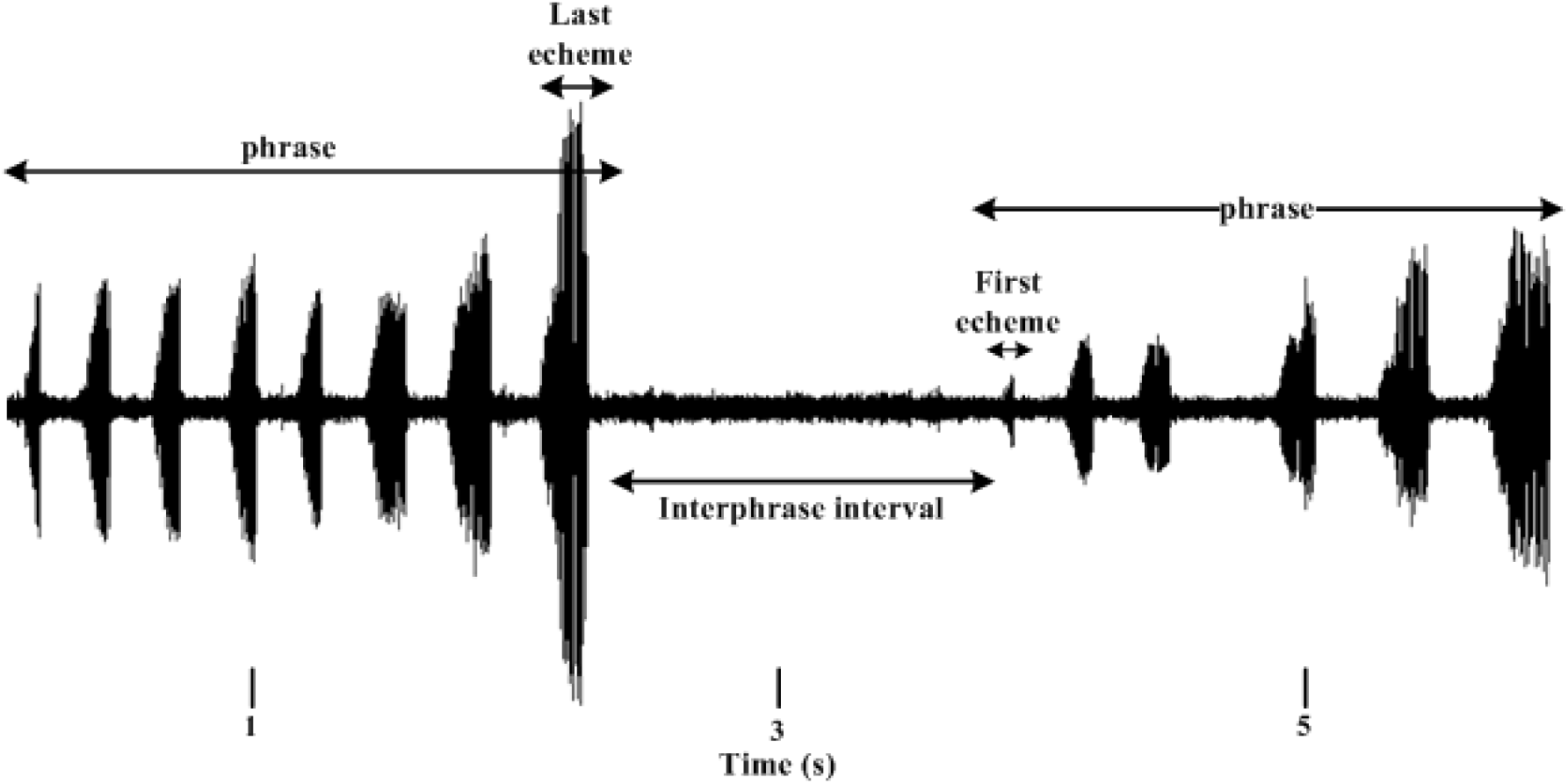
an example of echeme duration and amplitude increasing through the phrase: calling song of *Pagiphora annulata*

**Figure 3:**
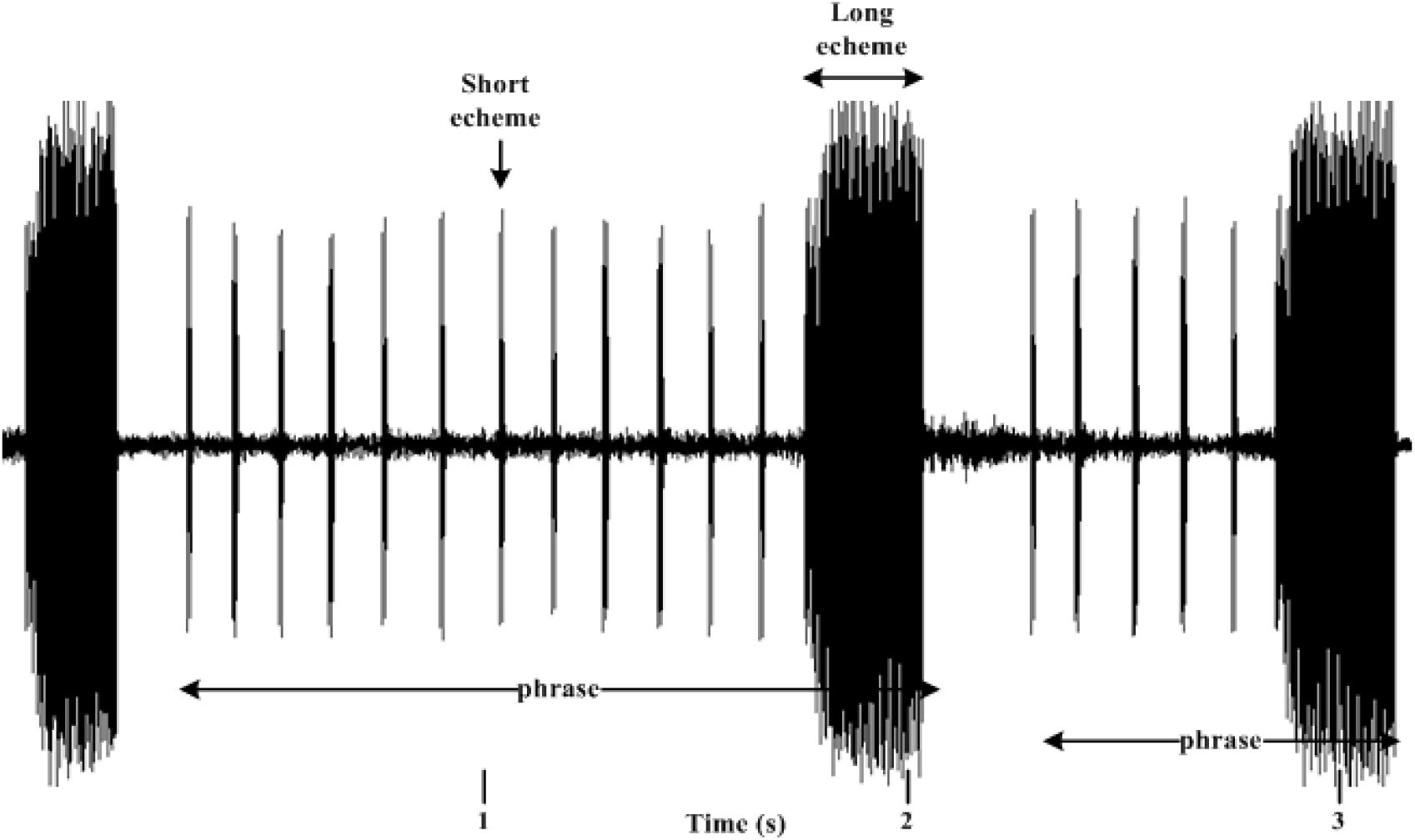
an example of Just two type of echeme; long echeme and short echeme: calling song of *Tettigetta golestani*

**Figure 4:**
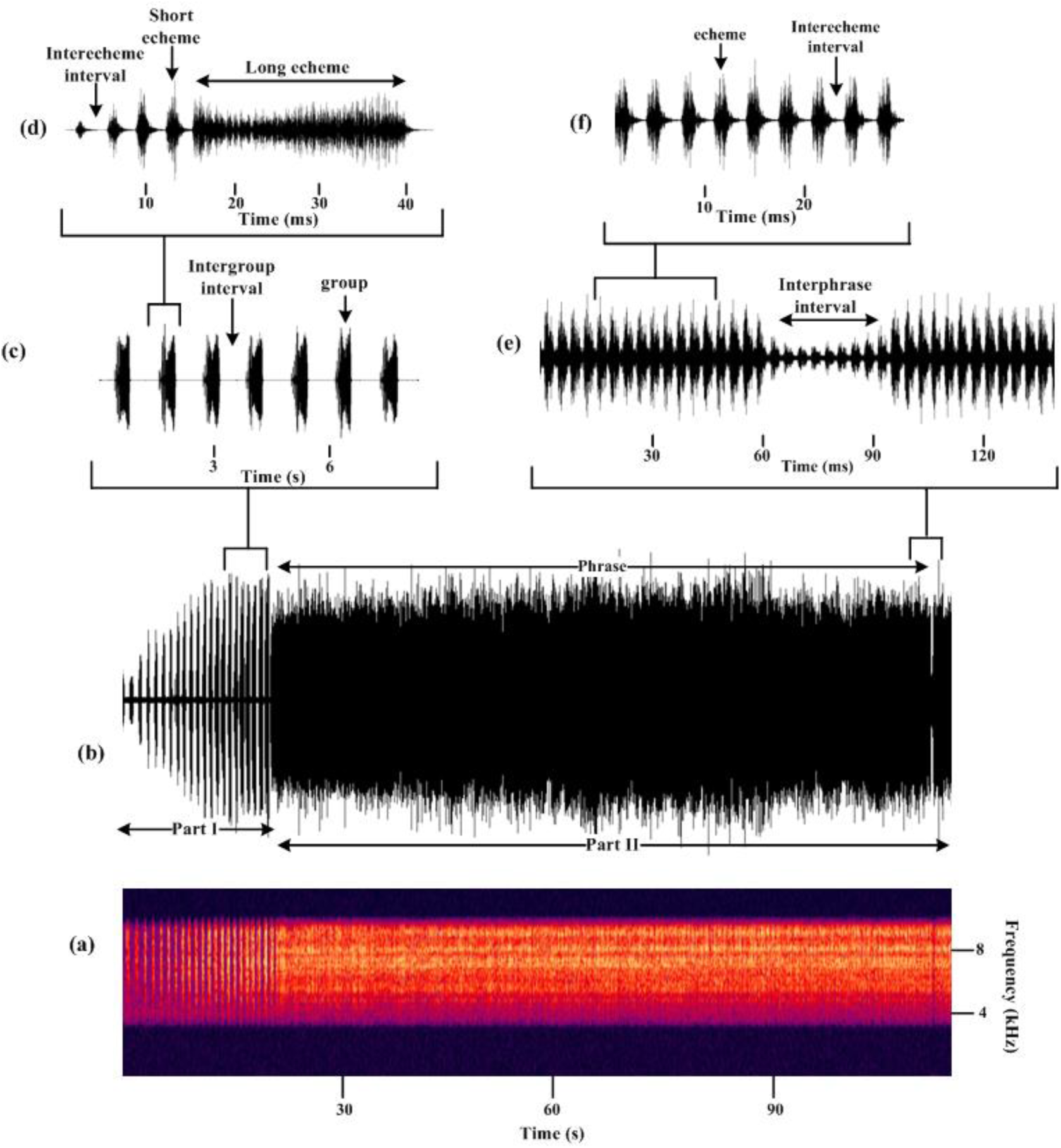
Calling song of *Chloropsalta smaragdula*; a: spectrum of the calling song; b: spectrum of calling song which is divided into two type part: part I and part II; c: part I consisted of some repeating groups and intervals between groups; d: subgroups of a group which is included several short and a long echemes (subgroup) and interecheme interval; e: Interphrase intervals which have very low amplitude and not stop or pause between them; f: Phrase of part II make up of repeating of echemes and interecheme intervals.

**Figure 5:**
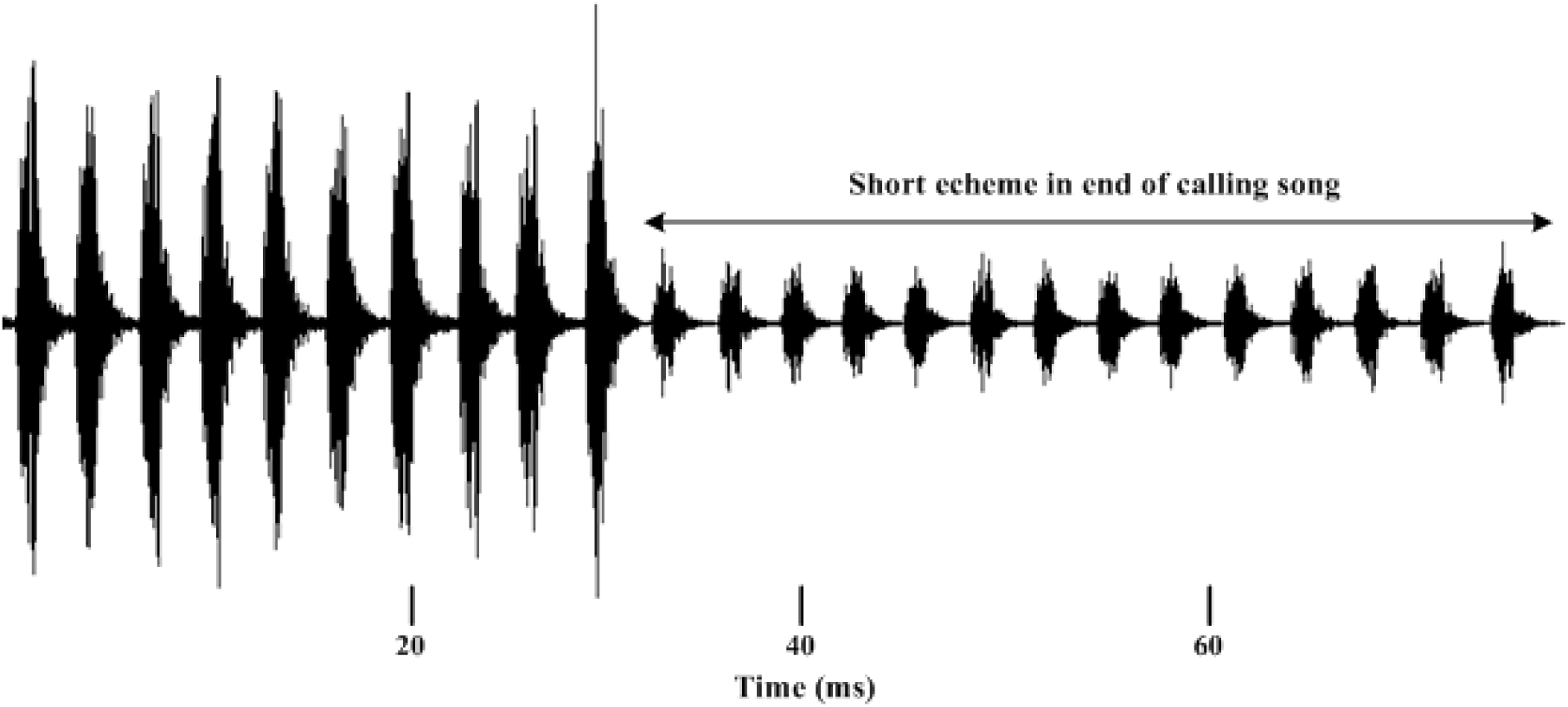
the end of calling song of one of six individuals of male of *Chloropsalta smaragdula* which is included of short echeme.

**Figure 6:**
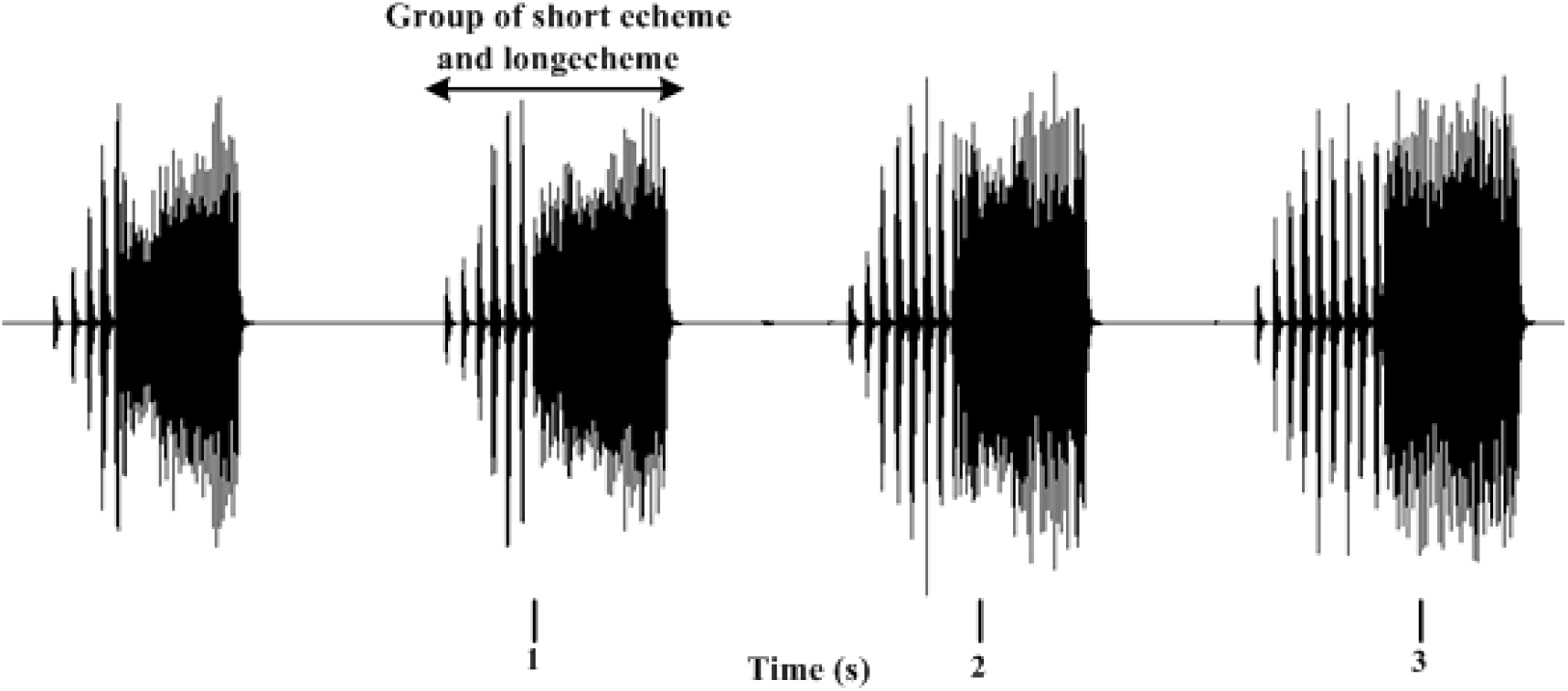
an example of phrase I including distinct groups of short and long echemes: calling song of *Chrysompalta smaragdula*

**Figure 7:**
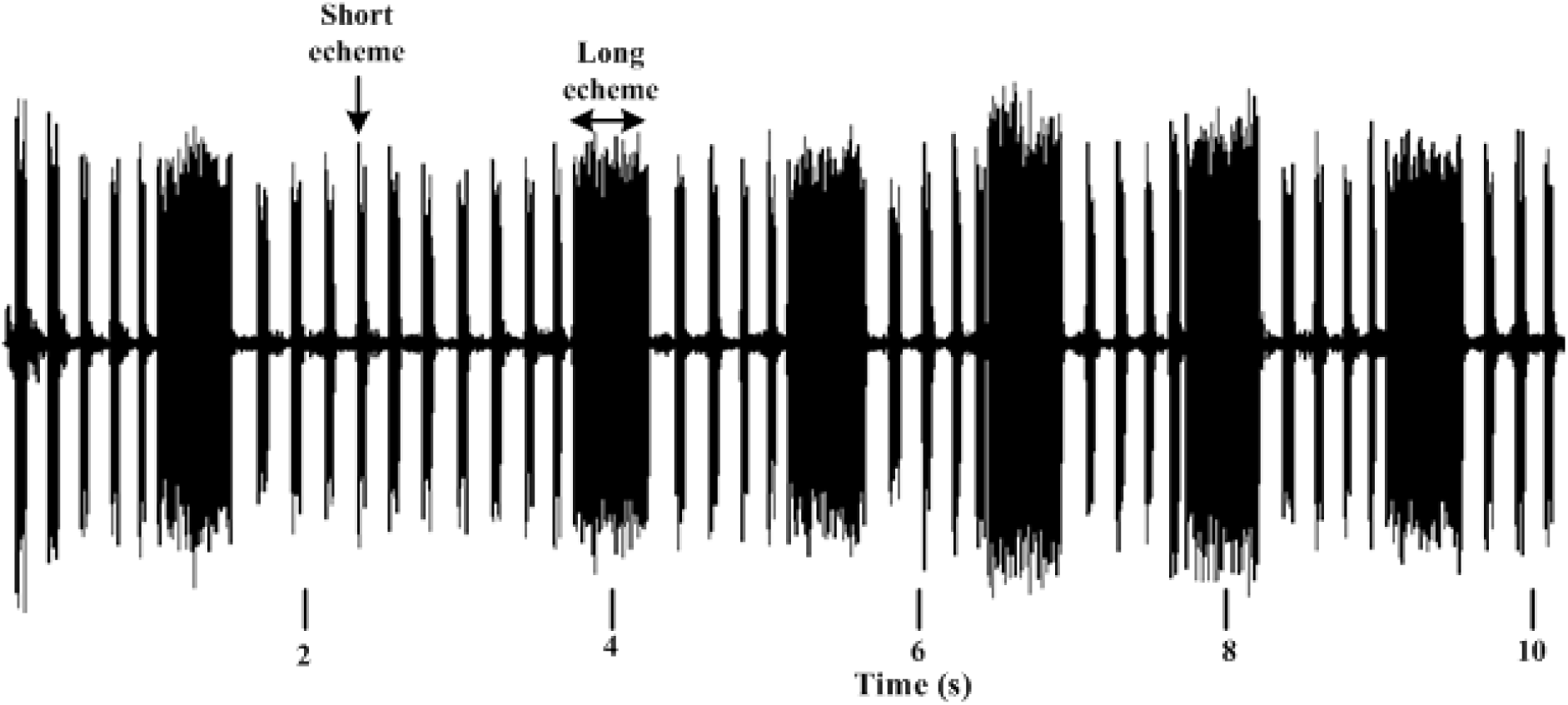
an example of phrase I including sequential short and long echemes: calling song of *Cicadivetta tibialis*

**Figure 8:**
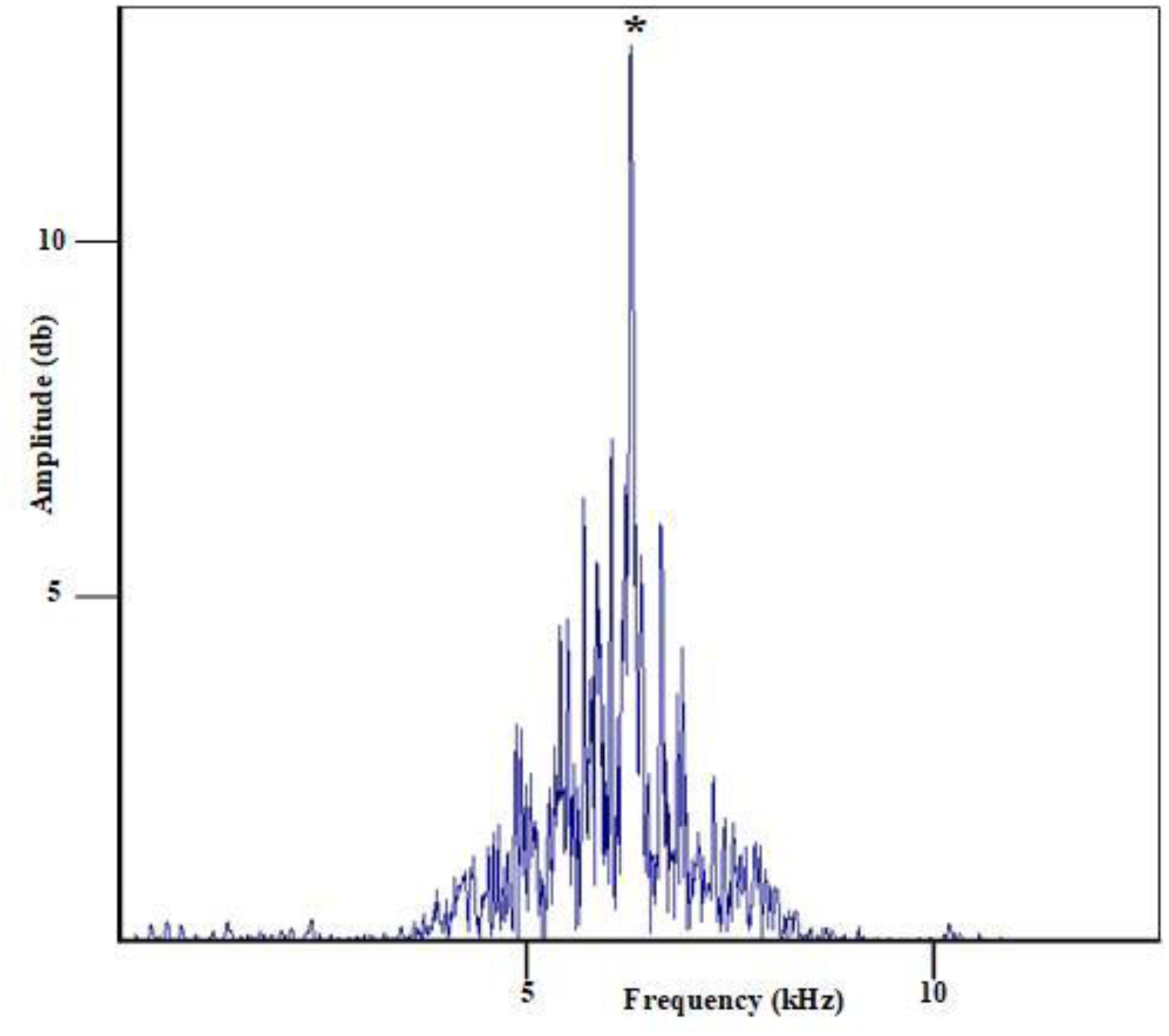
an example of one dominant frequency: dominant frequency of *Tibicen plebejus*: 6.12±0.44 kHz (Mehdipour et al., 2014)

**Figure 9:**
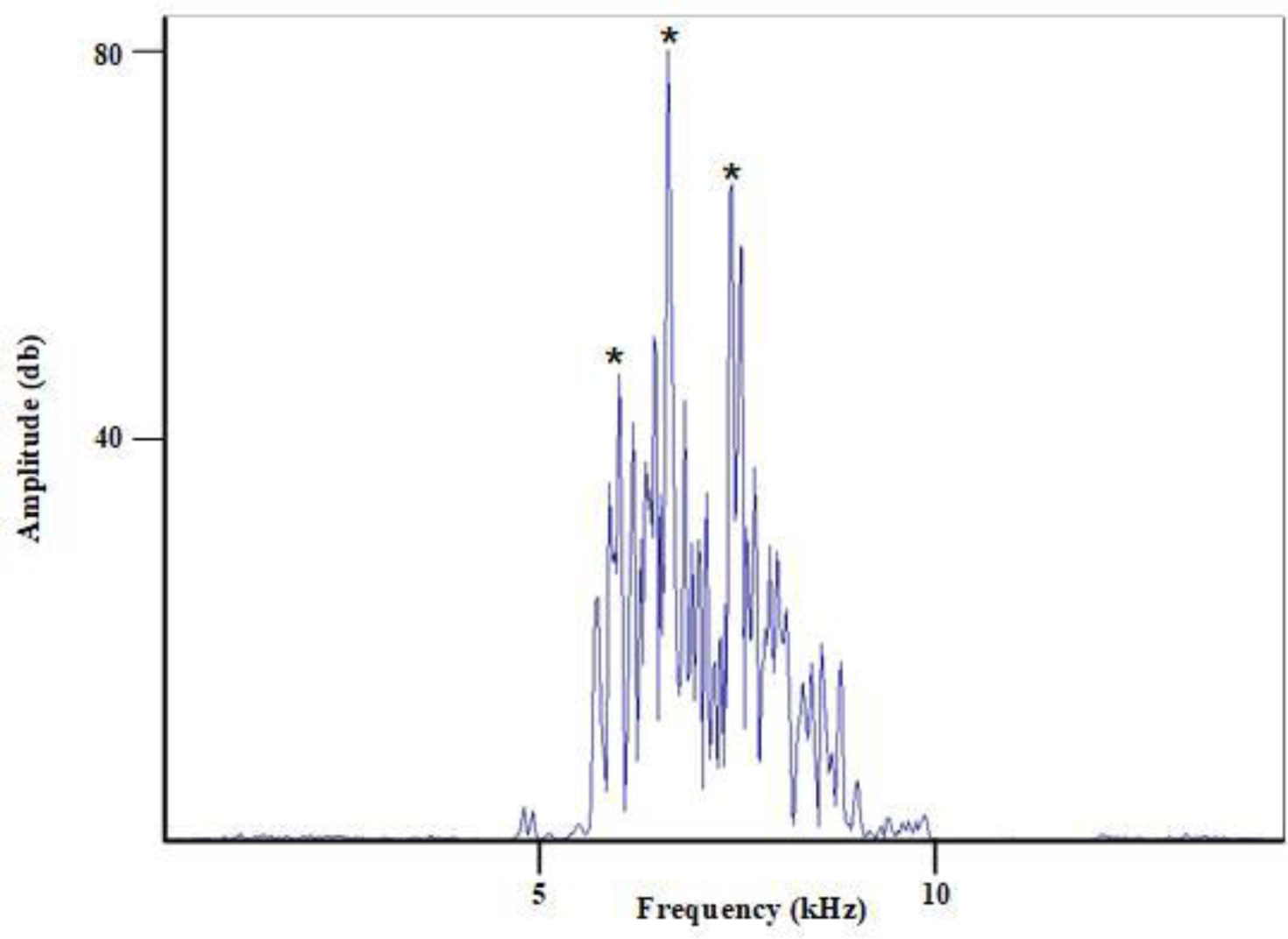
an example of more than one dominant frequency: dominant frequency of *Tibicina haematodes*: F1= 6.64±0.218 kHz, F2=7.51±0.247 kHz and F3=8.4±0.17 kHz (Sueur and Aubin, 2002; Sueur and Aubin, 2003)

In addition, here we introduce several alternative mathematical methods describing four complex spectral features (CSF) such as Variance, Kurtosis, Spectral centroid and Short time energy: The **variance** is the squared deviation from the mean of a random variable of amplitudes of segments of calling songs. The variance is also often defined as the square of the standard deviation. Variance is a measure of dispersion, meaning it is a measure of how far a set of amplitudes is spread out from their average value.

The **spectral centroid** is a measure that is used in digital signal processing to characterize a spectrum. The individual centroid of a spectral frame is defined as the average frequency weighted by amplitudes, divided by the sum of the amplitudes. The spectral centroid is used to find the center value of the groups for each insect frequency band. This feature is defined as,

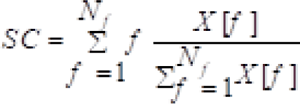

where the SC denotes the spectral centroid, f is the frequency domain, represents the normalized amplitude in the spectral domain and is the maximum frequency of each insect signal (Zamanian and Pourghassem, 2017).

**Kurtosis** is a measure of the tailedness of a distribution. Tailedness is how often outliers occur. Excess kurtosis is the tailedness of a distribution relative to a normal distribution. Distributions with medium kurtosis (medium tails) are mesokurtic. Distributions with low kurtosis (thin tails) are platykurtic. Datasets with low kurtosis tend to have a flat top near the mean rather than a sharp peak. A uniform distribution would be the extreme case which is defined as,

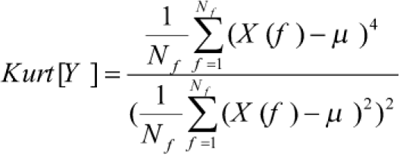

where is Kurtosis of the spectrum (Zamanian and Pourghassem 2017).

The **short time energy** is the energy of short signal (Yang et al. 2010). Short time energy is a simple and effective classifying parameter for all segments (Enqing et al. 2002). The short time energy of above signal can be determined from following expression:

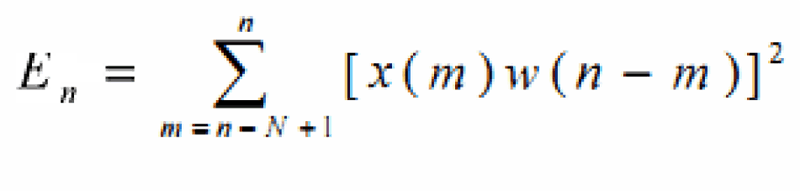

Where w (n-m) is the window, n is the sample that the analysis window is centered on, and N is the window length (Ubul et al. 2009).

CSFs are computed considering the spectrum as a distribution that the value of the distribution is replaced by the frequency of spectrum and the probability is replaced by the normalized amplitude. All of these features are extracted from the spectrum of signal which is considered as a distribution. The features are the statistical characteristics that does not matter which section of signal to analyze. They are extracted by MATLAB, and the code with data is available on github (https://github.com/honprogram/CicadaIran/)

### Calling songs used for re-analysis

Calling song recordings of *Cicadatra atra*, *Cicadatra persica* and *Psalmocharias querula* were provided by Matija Gogala and calling song of *Cicadatra barbodi* by Fariba Mozafarian.

The selected calling song of 17 and 60 seconds duration of *Cicadatra atra*, 40, 15 and 40 seconds of *Psalmocharias querula*, 65, 50 and 50 seconds of *Cicadatra persica*, 15 seconds of *Cicadatra barbodi* and 3 and 12 minutes of *Cicadatra alhageos* are sampled to 48 kHz by Cool Record Edit Deluxe Ver. 7.8.6. All of CSFs are listed in Table 1.

**Table 1:** four complicated spectral features of calling song such as Variance, Kurtosis, Spectral centroid and of *C. barbodi*, *C. atra*, *C. persica*, *C. alhageos* and *P. querula*.

| Feature<br>Cicada | Variance | Spectral centroid | Kurtosis | Short time energy |
| --- | --- | --- | --- | --- |
| *Cicadatra barbodi* | 80.14 | 12692.21 | 127.56 | 94.92 |
| *Cicadatra atra* | 51.13 | 20625.37 | 22.86 | 60.88 |
| *Cicadatra persica* | 32.90 | 16715.76 | 16.91 | 40.99 |
| *Cicadatra alhageos* | 22.27 | 9690.43 | 65.76 | 26.33 |
| *Psalmocharias querula* | 54.77 | 11306.79 | 33.02 | 60.62 |

### Recording and analysis of calling songs of *Chloropsalta smaragdula*

Research was conducted in Mobarakeh in Isfahan Province which is located in the west of Iran (51 ° 30’ eastern longitude and 32° 20’ northern latitudes; 1673 meters above sea level) in June 2011. In the natural environment in the vineyard, the calling songs of cicadas were recorded. A total of six males were collected. Recordings were performed with a digital Japanese sound recorder (SONY ICD-Sx713) with a sampling frequency range of 16 bits, and 44.1 kHz linear PCM, and a frequency response of 40 – 20,000 Hz. Calling songs of cicadas were analyzed by Cool Record Edit Deluxe Ver. 7.8.6 and Sound Ruler Ver. 0.9.6.0 on a desktop PC. Each calling song was recorded ten minutes. The microphone was placed at a distance of 30 cm from cicadas in the warmest hours of the day at noon when more cicadas were observed. Following variables of calling song of *Ch*. *smaragdula* were measured: Part I - Part I duration, number of groups, number of subgroup/groups, group duration, intergroup intervals, echeme duration, interecheme intervals; Part II: phrase duration, interphrase duration, echeme duration, interecheme interval, echeme period and dominant frequency. They were measured with a 0.001 s precision on signal oscillograms. Description of acoustic variables are explained in table 2.

**Table 2:** Descriptions of the calling song variables analyzed.

| Table 2: Descriptions of the calling song variables analyzed |  |  |
| --- | --- | --- |
| Variable |  | Description |
| part I | Rhythmic sound |  |
|  | Continuous sound | The regular repetition of a single echeme |
|  | Part I | Part I consists of some repeating groups in the beginning of calling song |
|  | Part I duration (s) | Duration of part I from start to end |
|  | Group | Group consists of short echemes and one long echeme in part I |
|  | No. of group | The number of group per part I |
|  | Group duration | Duration of each group from start to end |
|  | Intergroup interval | Interval between end of a group and beginning of next group |
|  | No. of echemes/<br>group | The number of short echemes per group |
|  | echeme | Each phrase consist of echemes basic units of sound (duration: hundredth of seconds or tenths of milliseconds) |
|  | Echeme duration | Duration of each echemes from start to end |
|  | Interecheme<br>interval | Interval between end of a echeme and beginning of next echeme |
| part II | Echeme duration | Duration of each echeme from start to end |
|  | Interecheme<br>"intervals" | Interval between end of a echeme and beginning of next echeme<br>here: parts with low amplitude of sound |
|  | Echeme period | Duration between the start of one echeme and the beginning of the<br>following one |
|  | phrase | Calling songs are repeated over long periods of time as called a<br>phrase |
|  | Phrase duration | Duration of each phrase from start to end |
|  | Interphrase interval | Interval between end of a phrase and beginning of next phrase |
|  | Dominant<br>frequency | The frequency of the maximum amplitude on the spectrogram |

## Results

This study describes a review of the calling song of 15 species of Iranian cicadas *Tibicen plebejus*, *Cicadatra lorestanica*, *Cicadivetta tibialis*, *Tettigetta golestani*, *Pagiphora annulata*, *Tibicina haematodes*, *Cicada orni, Pagiphora annulata*, *Chloropsalta smaragdula*, *Cicadatra hyalina*, *Psalmocharias querula*, *Cicadatra persica*, *Cicadatra alhageos*, *Cicadatra atra* and *Cicadatra barbodi* and their taxonomy via characteristics of calling songs. Based on standard and advanced parameters such as common acoustic methods, four new complex spectral features like variance, kurtosis, spectral centroid and short time energy, we present an acoustic identification key to theses Iranian cicadas. Ten species could be classified using VSTF extracted from different papers. However, determination of *Cicadatra atra*, *Cicadatra persica*, *Cicadatra barbodi*, *Cicadatra alhageos* and *Psalmocharias querula* required integration of CSF parameters into the key.

Our hierarchical identification system based on acoustic features was reliable and robust for species recognition. Terminology used in the key is either self-explanatory or described in the respective figures. For taxonomy we used the names of the respective bioacoustics’ publications cited here, which might deviate from the latest up-to-date taxonomy compiled by Dmitriev et al. 2027.

### Acoustic identification key to species of Iranian cicada

(1a) Rhythmic sound (Fig. 10) 2
(1b) Continuous sound (Fig. 11) 7
(2a) A single type of phrase 3
(2b) More than one type of phrase (Fig. 2) 5
(3a) A single type of echeme (Fig. 10) *Cicada orni*
(3b) More than one type of echeme 4
(4a) Echeme duration and amplitude increasing through the phrase (Fig. 3) *Pagiphora annulata*
(4b) Just two types of echeme; long echeme and short echeme (Fig. 6) *Tettigetta golestani*
(5a) Irregular echeme in phrase I *Cicadatra platyptera*
(5b) Regular echeme in phrase I 6
(6a) Phrase I including distinct groups of short and long echemes (Fig. 7) *Chloropsalta smaragdula*
(6b) Phrase I including sequential short and long echemes (Fig. 8) *Cicadivetta tibialis*
(7a) Calling song made up of echemes (Fig. 4e) 8
(7b) Calling song made up of pulses (Fig. 1) 9
(8a) One dominant frequency (Fig. 5) *Tibicen plebejus*
(8b) More than one dominant frequency (Fig. 9) *Tibicina haematodes*
(9a) One dominant frequency 10
(9b) More than one dominant frequency *Cicadatra hyalina*
(10a) Mean of dominant frequency≥ 11 kHz *Cicadatra lorestanica*
(10b) Mean of dominant frequency< 11 kHz 11
(11a) Mean of Kurtosis of calling song ≥ 100 *Cicadatra barbodi*
(11b) Mean of Kurtosis of calling song < 100 12
(12a) Mean of Spectral centroid of calling song ≥1900 *Cicadatra atra*
(12b) Mean of Spectral centroid of calling song <1900 13
(13a) Mean of Short time energy of calling song < 30 *Cicadatra alhageos*
(13b) Mean of Short time energy of calling song ≥ 30 14
(14a) Mean of Variance of calling song <40 *Cicadatra persica*
(14b) Mean of Variance of calling song ≥40 *Psalmocharias querula*

### Detailed description of calling songs

#### *Cicada orni* Linnaeus, 1758

Calling song pattern: Calling songs of *C. orni* is regular repetition of echemes with interecheme intervals, which constitute groups of impulses (Popov 1975, Claridge et al. 1979; Joermann and Schneider 1987; Fonseca 1991; Boulard 1995; Quartau et al. 1999; Quartau et al. 2000; Simões et al. 2000; Quartau and Simões, 2005; Pinto-Juma et al. 2005; Seabra et al. 2006; Simões et al. 2006; Sueur et al. 2010; Mehdipour 2011). Table 3 shows the time and spectral characteristics of the male cicada’s calling songs from different papers.

**Figure 10:**
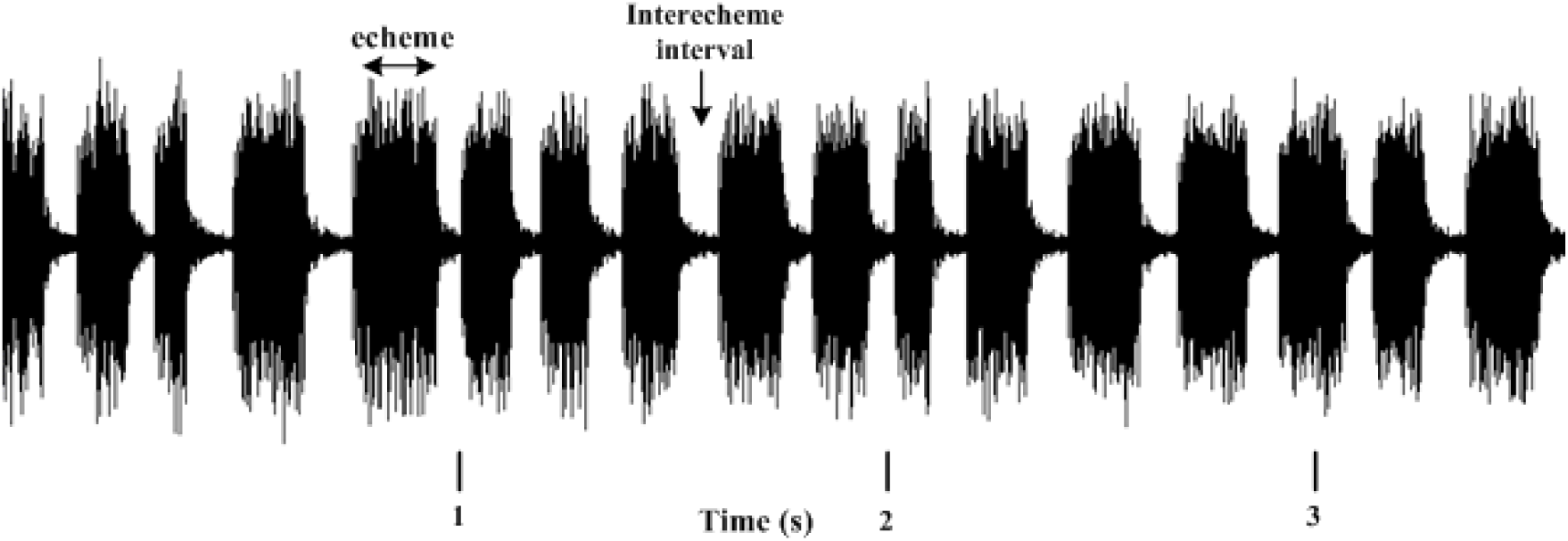
an example of rhythmic sound: calling song of *Cicada orni*

**Figure 11:**
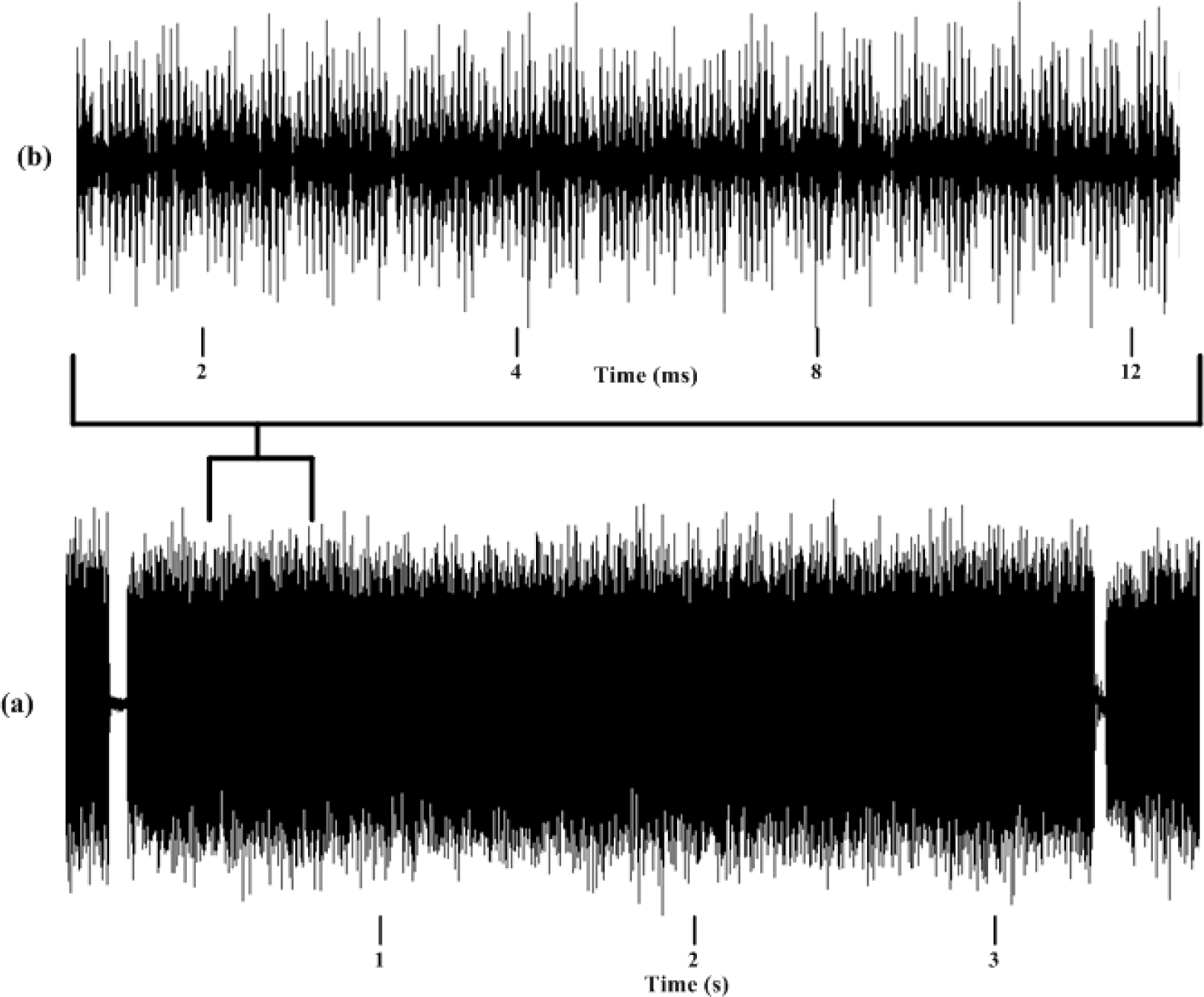
an example of continuous sound: (a) calling song of *Cicadatra alhageos*; (b) a part of calling song which consisted of pulses

**Table 3:** parameters of calling song of *Cicada orni* in different papers.

| Table 3: parameters of calling song of *Cicada orni* in different papers |  |  |  |  |  |  |
| --- | --- | --- | --- | --- | --- | --- |
| Variable<br>Author | Pinto-Juma *et al.* (2005) | Simões *et al.* (2000) | Sueur *et al.* (2010) | Quartau & Simões (2005) | Seabra *et al.* (2006) | Mehdipour (2011) |
| Echeme duration (s) | 0.08±0.03 | 0.093±0.03 |  | 0.07±0.02 | 0.078±0.03 | 0.07±0.01 |
| No. echeme/s | 4.69±1.30 | 2.923±0.66 |  | 4.15±1.13 | 5.37±1.33 | 5±1 |
| Interecheme interval (s) | 0.15±0.07 | 0.259±0.1 |  | 0.19±0.07 | 0.123±0.072 | 0.129±0.03 |
| Dominant frequency (kHz) |  |  | 4.51±0.15 |  |  | 4.914±0.33 |

Frequency parameter: The dominant frequency was reported around 4.51±0.15 kHz (Sueur et al., 2010) and 5.015±0.50 kHz (Simões et al., 2006) (Table 3).

#### Pagiphora annulata (Brullé, 1832)

Calling song pattern: The calling song consists of several phrases between 2.1 to 2.9 s, consisting of 7 to 10 echemes in each phrase. Phrases are separated by irregular intervals between 2 to 10 s. Repetition rate of echemes is 3±0.5 Hz (Gogala and Trilar 2000). Towards the end of each phrase, 3 to 5 echemes have longer duration and higher intensity due to additional wing clicking (Gogala and Trilar 2000; Gogala 2002).

Frequency parameter: The bandwidth is about 2.5 to 6 kHz (Gogala and Trilar 2000; Gogala et al. 2005) with dominant frequency of about 3.9 kHz (Gogala and Trilar 2000)

#### *Tettigetta golestani* Gogala and Schedl 2008

Calling song pattern: Gogala and Schedl (2008) reported that the calling song comprises repeating phrases with two kinds of echeme: a short echeme of 11.5±2.1 ms and a long echeme at the end of each phrase, with length of 183±42 ms. Frequency parameter: The dominant frequency is 10.5 kHz (Gogala and Schedl 2008).

#### Cicadatra platyptera Fieber 1876

Calling song pattern: There are irregular echemes in the beginning of calling song (ranging from 5–15 ms) but in the middle of calling song, echeme duration is 122.7±16.04 (ranging from 50–188 ms) and interecheme intervals are 91.2±17.63 (ranging from 40–213 ms). If *C. platyptera* produce calling song after courtship which did not lead to mating, echeme duration of calling song is slightly different, 133±42.22 (ranges of 75–277 ms) and interecheme intervals are 80±18.65 (ranges of 36–212 ms) (Mol et al. 2013).

Frequency parameter: The bandwidth of calling song is from about 5.5 to 12 kHz with a maximum 6, 8 and 10 kHz (Mol et al. 2013)

#### Chloropsalta smaragdula Haupt 1920

This species is known from Turkey, Iraq and Iran (Metcalf 1963; Dlabola 1981; Duffels and Van der Laan 1985; Mirzayans 1995). Here we present recordings and describe calling song parameters of this widely distributed species for the first time. Notably, *Ch*. *smaragdula* sings in a head-downward position.

Calling song pattern: The calling song can last for some minutes, without interruption. It consists of two parts. Part I is observed at the beginning of the calling song (Fig 4a,b). Part I consists of repeating subgroups of several short and long echemes, separated by intervals (Fig4c). In subgroups, the last echeme is longer than other echemes but the last interecheme interval is shorter than others (Fig 4d). The number of subgroups in each cicada and the number of echemes in each group is of 31.75±15.04 and 6.28±1.33 (mean±stdev) respectively.

Part II follows directly and consists of irregular repetition of echemes separated by irregular intervals (Fig 4b). These “intervals” between phrases are not silent but consist of echemes of very low amplitude, with no real pause between them (Fig. 4e). Phrases are composed of regularly repeated echemes and interecheme intervals (Fig. 1f). The end of calling song of just one from six individuals was with several regular repeated short echemes (Fig. 9). Table 4 shows the temporal and spectral characteristics of the male cicada’s calling songs.

**Table 4:**
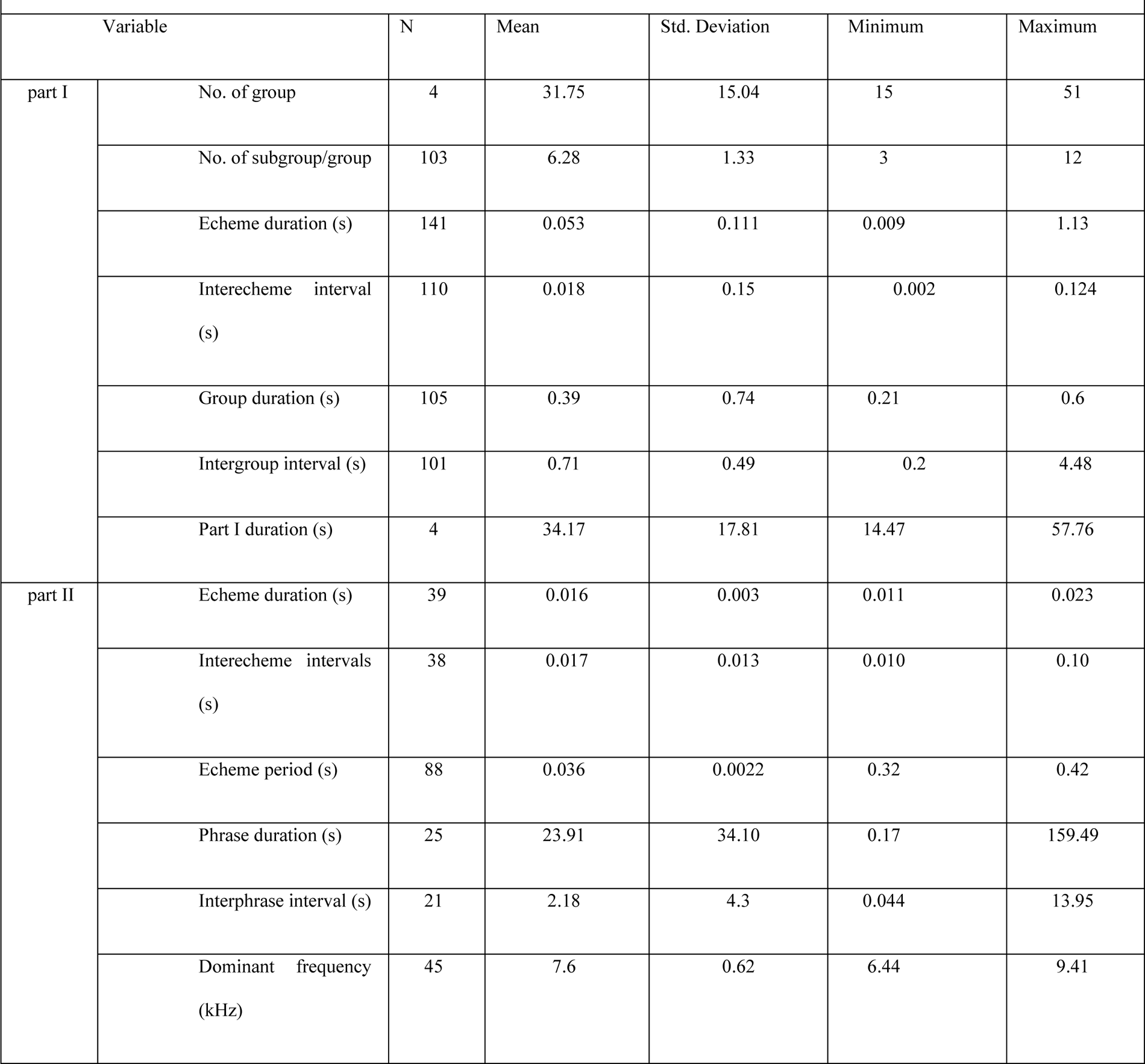
parameters of calling song for *Chloropsalta smaragdula* males. N=Sample size.

Frequency parameter: Dominant frequency band from 6.44 to 9.41 kHz (mean 7.6±0.62 kHz).

#### Cicadivetta tibialis (Cicadetta tibialis) (Panzer 1798)

Calling song pattern: The rhythmic calling song can be seen in *C. tibialis* (Boulard 1995; Gogala et al. 1996; Puissant and Sueur 2010). The calling song repertoire of *C. tibialis* seems to contain two types of phrases and echeme or binary sound (Sueur and Puissant, 2000; Gogala 2002; Puissant and Sueur, 2010). Phrase I (beginning of calling song) has a long echeme (307±59 ms) with series of short echemes (51±10 ms) and the last short echeme (39±7 ms) (Sueur and Puissant 2000) is shorter than other short echemes (Gogala et al. 1996; Sueur and Puissant 2000) and duration of phrase I is 1025±508 ms (Sueur and Puissant 2000). The number of short echemes varies in each phrase I between 2 and 15 according to Gogala et al. (1996), but Sueur and Puissant (2000) report 4.8±2.8. The first interval between long echeme and short echeme is shorter (103±19 ms by Gogala et al. (1996) and 102±15 ms by Sueur and Puissant (2000)) than subsequent intervals between short echemes (119±26 ms by Gogala et al. (1996) and 118±15 ms by Sueur and Puissant (2000)). The interval between last short echeme and next long echeme is 69±20 ms (Sueur and Puissant 2000)

Phrase II (calling song) consists of regularly repeated short echemes (59±7 ms). Interval between echemes of phrase II (172±25 ms) (Sueur and Puissant 2000) is longer than those between the short echemes of phrase I (Gogala et al.; Sueur and Puissant 2000).

Frequency parameter: 2 bands of frequency spectrum can be seen, a principal one between 12-22 kHz with a maximum between 14 – 18 kHz, and. a second band between 7-8 kHz (Gogala et al. 1996).

#### Tibicen plebejus (Scopoli 1763)

Calling song pattern: the calling song includes repeating phrases of some echemes that last without interruption for some minutes (Claridge et al. 1979; Joermann and Schneider 1987; Boulard 1995; Gogala 2002; Sueur et al. 2004; Puissant 2012; Mehdipour 2011; Mehdipour et al. 2014). Phrase period 7-30 s according to Joermann and Schneider (1987), coinciding with 6.25-33.93s by Mehdipour et al. (2014). A phrase consists of three parts with distinct amplitudes (Claridge et al. 1979; Joermann and Schneider 1987), or two parts according to echemes (Mehdipour et al. 2014). As far as amplitude is concerned, (1) signal begins from low amplitude and quickly increases to maximal amplitude, (2) amplitude remains maximal and (3) amplitude decreases. With respect to echemes; (1) phrase part 1 (10.31±3.74 s): signal has echemes (echeme period 80.15±7.01 ms and echeme duration 54.57±7.63 ms) and interecheme interval (25.82±9.83 ms) is the same as first and second part of phrase that is described by Claridge et al. (1979). Joermann and Schneider (1987) calculated echeme period 84.6±2ms. (2) Phrase part2 (1.86±0.67 s): The amplitude of phrase 2 is low. The low amplitude characteristic of phrase2 is similar to the third part of phrase by Claridge et al. (1979) (Mehdipour et al. 2014). The number of phrase/min 5.29±1.03, the number of echeme/s 12.24±0.89, echeme/interecheme ratio 2.31±0.91s (Mehdipour et al. 2014).

Frequency parameters: Dominant frequency is calculated 5.48±0.77 kHz by Sueur et al. (2010), 5.077±0.2kHz by Mehdipour (2011), around 6-7 kHz by Puissant (2012) and 6.12±0.44 kHz by Mehdipour et al. (2014). The bandwidth ranged between 4 to 8 kHz (Mehdipour et al. 2014).

#### Tibicina haematodes (Scopoli, 1763)

Calling song pattern: The signal is composed of a succession of phrases (around 14.0± 2.3 ms) consisting of two parts. First part is the beginning of calling song and second part is the main calling song. The first part consists of about 4 successive short echemes, and the second part consists of echemes and interecheme intervals (echeme duration 8.2±0.4 ms; echeme period 10.2±0.4 ms; the number of echeme per second 98.3±1.6) that consists of two subgroups of pulses. The pulses last approximately 1 ms. Each echeme is made up of 6-8 pulses arranged in two subgroups containing 3-4 pulses (Sueur and Aubin 2002; Sueur and Aubin 2003).

Frequency parameter: The frequency spectrum has three main peaks: F1= 6.64±0.218 kHz, F2=7.51±0.247 kHz and F3=8.4±0.17 kHz (Sueur and Aubin 2002; Sueur and Aubin 2003).

#### Cicadatra hyalina (Fabricius, 1798)

The calling song of *C. hyalina* involved as continuous signal and echeme song. Also, this species has two types of phrases, a continuous one and a regularly repeated phrase with “rumbling sound” (Popov 1975; Boulard 1995). Duration of continuous song is, on average, 1.3-9 minutes or longer. A continuous signal consists of regular syllables which are made of 5-6 pulses, lasting, on average, 5-6 ms. After some minutes, this continuous signal changes into a short echeme song called rumbling sound by Boulard et al. 2018. It is repeated regularly, lasting between 640-8952±651ms. Echemes are separated by intervals between 232-704±149 ms. Echemes consist of two connected pulses of 2-3 ms length, separated by intervals between 14-16 ms (Mol 2018).

Frequency parameters of the calling song are characerized by a wide range with distinct peaks (Popov 1975; Boulard 1995). Mol (2018) measured a frequency range from from 10248 to 16760 Hz, with maxima at 10248 Hz, 11791 Hz and 15163 Hz.

#### *Cicadatra lorestanica* Mozaffarian and Sanborn, 2010

Calling song pattern: The calling song of *C. lorestanica* is continuous and consists of phrases of irregular duration. There is a continuous series of pulses (Mozaffarian et al. 2010; Mehdipour 2011). The repetition rate of pulses per second is 917±68 (Mozaffarian et al. 2010).

Frequency parameters: Dominant frequency is 11.39±0.09 kHz according to Mozaffarian et al. 2010, and 11.886±0.776 kHz according to Mehdipour (2011).

#### *Cicadatra barbodi* Mozaffarian and Sanborn, 2013

Calling song pattern: *C. barbodi* calling song consists of a series of short and long phrases (in average 1.442s ranging from 0.0661 to 2.644), with short intervals between phrases (in average 0.047 s ranging from 0.033-0.057 ms). Each phrase is composed of a continuous series of pulses (0.001 ms length). On average, 1.5 phrases per second were generated (Mozaffarian and Sanborn 2013).

Frequency parameters: The dominant frequency is 9.562 kHz (Mozaffarian and Sanborn 2013).

#### Cicadatra atra (Olivier 1790)

Calling song pattern: The calling song of *Cicadatra atra* is recorded as a continuous song (song or phrase type 1) with irregular interruptions and a series of short echemes (song or phrase type 2) (Gogala and Trilar 1998; Sueur et al. 2006; Mol et al. 2013) The repetition rate of echemes in the phrase type 2 is 2.5 per second (Simões et al. 2013).

Frequency parameters: The dominant frequency is described at 10.23±0.78 kHz by Simões et al. (2013) and 9.8 kHz by and Boulard (1992). The bandwidth is between 6 and 17 kHz (Sueur et al. 2006).

#### Cicadatra alhageos (Kolenati 1857)

Calling song pattern: The calling song consists of a repeating phrase (0.188 to 15.43 s) with irregular interruptions (6 to 623 ms). At the beginning of calling song, there are some irregular echemes (7 to 773 ms) with interecheme intervals (4 to 567 ms) (Mehdipour et al. 2015). Zamanian et al. (2008) differentiates two sections: ‘start sound’ and ‘continued sound’. The irregular echemes during the first seconds of the calling song belong to the start sound section. The continued sound section includes phrase and inter-phrase intervals.

Frequency parameter: Dominant frequency is recorded from 10.2 kHz and 9.47±1.2 kHz and 9.27 to 10.5 kHz (mean 10.1±0.28 kHz) by Zamanian et al. (2008), Mehdipour (2011) and Mehdipour et al. (2015), respectively and frequency bandwidth is 6 kHz (Mehdipour et al. 2015).

#### Cicadatra persica Kirkaldy, 1909

Calling song pattern: Calling song of *C. persica* is continuous, with phrases uninterrupted for many minutes (Gogala and Trilar 1998; Dadar et al. 2013). Each phrase consists of a continuous series of pulses, with one or a few irregular echemes towards the end (Gogala and Trilar 1998).

Frequency parameters: bandwidth is recorded between 5.8 and 12.4 kHz with dominant frequency near 8.4 kHz, according to Gogala and Trilar (1998) and also Dadar et al. (2013) described that the frequency ranged between 1.051.8 ±0.041 kHz and 5.74 ± 0.010 kHz with frequency maximum amplitude between 2.72 ±0.0 24 kHz and 4.70 ±0.0 52 kHz by. The beginning of a continuous song is without any clear pattern of amplitude modulation (Gogala and Trilar 1998).

#### Psalmocharias querula (Pallas, 1773)

Calling song pattern: The song of *P. querula* is a continuous buzzing (song 1) and a series of shorter sequences combined from a short high pitched echemes and rumbling sounds (song 2) (Popov 1975; Gogala 2009).

Frequency parameter: Peak frequency of 5–7 kHz and phrases of calling song are amplitude and frequency modulated at the beginning of each phrase of the song 2 (Popov 1975).

## Discussion

We demonstrated that acoustic features of cicada songs can be used in cicada taxonomy for species recognition with high accuracy. Our taxonomic key relies on a hierarchical arrangement of simple visible spectral and temporal signal features (VSTF) which are sufficient to identify 10 species. To identify the remaining species, we showed that four complex spectral features (CSFs) were sufficient. We conclude that CSFs provide valuable additional bioacoustic parameters, and we provide open access code using MATLAB tools via github.

In the large number of cicadas, it is more complicated to use only VSTF to produce a key. So, classifying cicada based on physical characteristics of calling song is some cases not enough. In this case, entomologists must know other type of acoustic signals such as courtship songs which maybe do not contribute to complete acoustical taxonomy. It is better to add CSF being a useful method even if all types of acoustic signals are unavailable.

Some studies have been conducted on acoustical taxonomy of singing insects. Two groups of singing insects; crickets and cicadas are categorized by non-parametric (probabilistic neural network) and parametric (Gaussian mixture models) estimation of the probability density function with accuracy that exceeds 99% on the levels of family, and 90% on the level of species (Potamitis et al. 2007). Employing Melfrequency Cepstrum Coefficient (MFCC) as sound feature demonstrates the automatic acoustical identification’s accuracy of above 96% (Zhu 2011). Noda et al. (2016) describe those signals have been characterized by MFCC and also Linear Frequency Cepstral Coefficient (LFCC) which shows high classification rates of 99.08%, surpassing other methods based on probabilistic algorithms. Other methods such as temporal and spectral features identify cicadas with accuracy of 98% (Zamanian and Pourghassem 2017). These researchers and others show that acoustical identification by a variety of methods can help entomologists to develop acoustical classification of singing insects with high accuracy.

Our approach can easily be combined with more sophisticated recognition software, which is mainly based on neural networks and already works sufficiently well for image-based plant identification (e.g. flora incognita app: https://play.google.com/store/apps/details?id=com.floraincognita.app.floraincognita&hl=de&gl=US) and birdsong recognition (https://birdnet.cornell.edu/). However, the underlying algorithms require considerable computer power, and therefore are server-based, sending results to users who uploaded multimedia data via apps. Hierarchical recognition tools as presented here could be combined with such server-based AI tools and apps in an interactive fashion. This would also help to aggregate more acoustic data from citizens. Nowadays, high quality recording equipment is cheap and small. Even smartphone recordings could contribute to collect songs of species from a wide variety of habitats, from suburbs to jungles. Citizen science platforms such as naturalist (https://www.inaturalist.org/) and xenocanto (https://xeno-canto.org/) already allow upload of acoustic files, replay and sonagram visualisation (xenocanto), but hitherto do not provide recognition tools. Some server-based simple automatized tools such as extraction of carrier frequency or even CSF features might be a feasible intermediate step to help users identify their recordings.

In summary, recording insect songs is a cheap and reliable a valuable method for taxonomy, systematics, and biological monitoring.

Unfortunately, up to now only very few cicada reference recordings are publicly available We therefore want to encourage researchers to cooperate and share their acoustic recordings, to accelerate the development of computer-aided recognition tools.

## Acknowledgements

We are grateful to Matija Gogala for continual support and also for recordings of *T. golestani*, *Psalmocharias querula, Cicadatra atra* and *Cicadatra persica*. We also thank Fariba Mozaffarian to send recordings of *C. barbodi*.

